# A reinforcement learning algorithm shapes maternal care in mice

**DOI:** 10.1101/2022.03.21.485130

**Authors:** Yunyao Xie, Longwen Huang, Alberto Corona, Alexa H. Pagliaro, Stephen D. Shea

## Abstract

The neural substrates for processing classical rewards such as food or drugs of abuse are well-understood. In contrast, the mechanisms by which organisms perceive social contact as rewarding and subsequently modify their interactions are unclear. Here we tracked the gradual emergence of a repetitive and highly-stereotyped parental behavior and show that trial-by-trial performance correlates with the history of midbrain dopamine (DA) neuron activity. We used a novel behavior paradigm to manipulate the subject’s expectation of imminent pup contact and show that DA signals conform to reward prediction error, a fundamental component of reinforcement learning (RL). Finally, closed-loop optogenetic inactivation of DA neurons at the onset of pup contact dramatically slowed emergence of parental care. We conclude that this prosocial behavior is shaped by an RL mechanism in which social contact itself is the primary reward.

**One-Sentence Summary:** Maternal interactions with offspring are shaped by a dopaminergic reinforcement learning mechanism.

---

The ability to process and respond to rewards is a fundamental aspect of brain function. A confluence of data and theory supports a framework for understanding how organisms adapt their behavior to maximize reward known as ‘reinforcement learning’ (RL) (*1, 2*). Two crucial features of most RL models are iteration and reward prediction. Over many trials, an agent selects actions predicted to bring reward, and on each trial it compares the received reward with the predicted reward. The difference, called ‘reward prediction error’ (RPE), is then used to update subsequent actions and reward predictions. There is a wealth of evidence that RPE is explicitly signaled by midbrain dopamine (DA) neurons in response to rewards (*3, 4*).

For many animals, including humans, social contact is highly rewarding (*5*). Nevertheless, how social rewards are encoded in the brain and used to modify social behavior is poorly understood. Recent studies revealed some of the neural substrates for regulating sociality and sensing social reward (*6–11*), but the temporally unstructured nature of most social interactions present a challenge for observing their cumulative reinforcing influence on behavior. Here we use the gradual emergence and refinement of maternal care in virgin female mice to examine the relationship between midbrain DA reward signals and performance over many iterations of a stereotyped ethological behavior.

When first exposed to offspring, over several days, primiparous and virgin female mice exhibit increasingly rapid and reliable retrieval of pups that become separated from the nest (*12, 13*). We chose to examine the role of midbrain dopamine signals in the emergence of maternal retrieval for several reasons. First, access to pups positively reinforces behavior with comparable efficacy to cocaine (*14–17*). Second, the dopaminergic system has been linked to the establishment and motivation of pup retrieval (*18–23*). Third, the repetitive, stereotyped nature of retrieval facilitated a trial-by-trial analysis of DA signals and performance.

To initially observe DA neuron activity during pup retrieval, we injected the ventral tegmental area (VTA) of virgin female DAT-Cre^+/−^ mice with Cre-dependent adeno-associated virus (AAV) driving expression of jGCaMP7f (*24*), a genetically-encoded Ca^2+^ sensor (Fig. 1A). This resulted in expression of GCaMP in VTA DA neurons with high sensitivity and specificity (Fig. 1B). We then used fluorescence fiber photometry to measure fluctuations in population neural activity during maternal interactions.

**Figure 1:**
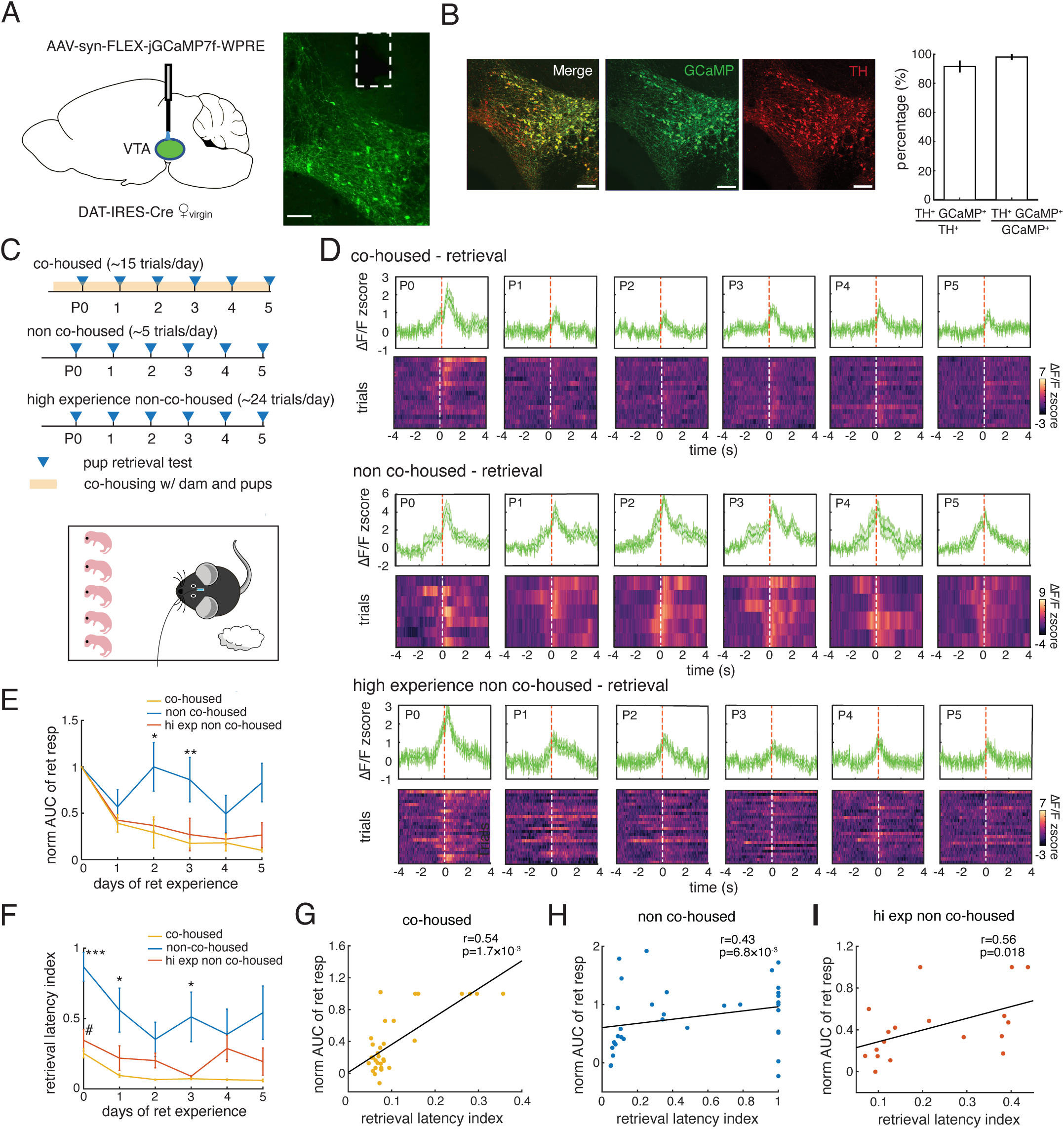
Population activity of VTA DA neurons during maternal retrieval is inversely related to retrieval performance. **(A)** Intersectional viral expression strategy for fiber photometry. **(B)** Co-localization of GCaMP7f and VTA DA neurons. Scale bar=100 µm. **(C)** Schematic of experimental design for retrieval tests comparing the effects of co-housing and practice on retrieval performance and VTA activity. **(D)** Dynamics of VTA DA neuron responses for P0 – P5 in three cohorts. Each row of plots depicts daily retrieval responses for one mouse with a heatmap of activity from all trials (*bottom*) and a trace of mean responses for each day (*top*, green trace). Each row of plots consists of data from a representative mouse from one of three cohorts: ‘Co-housed’, ‘Non Co-housed’, and ‘High Experience Non Co-housed’. See main text for descriptions of cohorts. **(E)** Plot of mean ± SEM magnitude of fluorescent signals over all experimental days separated by cohort. Magnitude is quantified as the mean integrated area under the curve (AUC) of the Z-scored response trace, normalized to the value on P0 (n=6, 7 and 3 for co-housed, non co-housed and hi exp non co-housed mice respectively, two-way ANOVA with Tukey correction, main factor (day): *p* < 0.001, main factor (mouse type): *p<0*.*0001*; * represents significant difference between the CH and the NCH groups, \**p* < 0.05, \*\**p* < 0.01). **(F)** Plot of mean ± SEM of retrieval latency index over all experimental days separated by cohort (same numbers of mice for each group as in E, two-way ANOVA with Tukey correction, main factor (day): *p>0*.*05*, main factor (mouse type): *p<0*.*0001*; * and # represent significant difference in the CH vs the NCH groups, and in the CH vs HE/NCH respectively, \**p* < 0.05, \*\**p* < 0.01, \*\*\**p* < 0.0001). **(G – I)** Scatterplots and Spearman correlation of normalized AUC and retrieval latency for ‘Co-housed’ (G), ‘Non Co-housed’ (H), and ‘High Experience Non Co-housed’ (I) cohorts.

When we performed daily pup retrieval tests (P0 – P5) on virgin mice while continuously co-housed (CH) with a new mother and her pups (Fig. 1, C and D), we observed large, brief transients that began at pup contact (Fig. S1) and peaked just after the female lifted the pup (Supplementary Video 1). Retrieval-evoked VTA activity was initially of comparable magnitude to responses to consumption of an unexpected treat (Fig. S2). Signals were largest on P0, and on subsequent days, the amplitude decreased sharply (Fig. 1, D and E).

To assess whether the decrease in DA neuron signals was due to continuous exposure to pups, we tested retrieval in a separate ‘Non co-housed’ (NCH) cohort that only encountered pups during one daily round of testing (5 trials) and then remained in their home cage with other virgins only (Fig. 1, C and D;). In contrast to CH mice, NCH subjects exhibited large amplitude GCaMP signals that were sustained through the end of the experiment (P5) (Fig. 1, D and E).

Finally, we reasoned that the differences in VTA activity and behavioral performance between the CH and NCH groups could directly reflect housing conditions, or simply the additional interaction time experienced by the CH group. We therefore performed retrieval tests on a third cohort we refer to as ‘High experience, non co-housed’ (HE/NCH; Fig. 1, C and D). These mice were housed separately from the dam and pups, but they were given many more trials per day (~24). Their DA neuron activity diminished rapidly after P0 like the co-housed group (Fig. 1, D and E), showing that the decline was related to the quantity of practice the mice had and not housing condition. ‘

In all three groups, the changes in DA signals were related to the strength and pace of learning, as measured by improvement in retrieval latency index (see Methods). The CH and HE/NCH mice showed indistinguishably robust and rapid learning (Fig. 1F). In contrast, the persistent, high-amplitude signals in NCH were accompanied by poorer retrieval (Fig. 1F). Learning in all cohorts was closely related to VTA activity because performance on each day was inversely correlated with response magnitude (Fig 1, G to I).

Our observation that responses to pup contact resembled responses to treats suggest that contact with pups may be intrinsically rewarding. RPE is a computed quantity that is commonly signaled by VTA. Because RPE is the difference between expected reward and received reward, as a reward becomes predictable, VTA responses to it diminish. We speculated that the decrease in VTA responses to pups may result from VTA encoding an RPE for pup contact. If so, then it should exhibit three properties. First, it should incrementally influence retrieval behavior on a trial-by-trial basis during early maternal experience. Second, surprising reward should evoke a larger RPE, while surprising lack of reward should evoke a negative signal. Third, as reward becomes increasingly predictable due to a reliable cue, RPE should decrease for the reward and increase for the predictive cue.

As the virgin mice accrued varying levels of maternal experience, retrieval time and mean retrieval-related DA neuron activity steadily decreased. However, both quantities showed considerable trial-to-trial variability (Fig. 2A). Therefore, we more closely examined the relationship between behavior and the recent history of VTA responses to pups. We used the velocity of the female’s approach to the pup as a single-trial measure of behavioral motivation and performance. Like latency index, mean approach velocity steadily improved over days of training (Fig. 2B). The higher velocity in the HE/NCH group may reflect the fact that those mice participated in more retrieval trials than the other groups (see Methods). As expected, for all three groups, the approach velocity on each trial was inversely correlated with the VTA signal on the same trial (Fig. 2, C and E and G). This was not true for adjacent trials. The difference in approach velocity from the previous trial was significantly positively correlated with VTA activity on the previous trial (Fig. 2, D and F and H). These correlations were abolished by shuffling trials (Fig. 2I). This result implies that the mouse updates its behavior trial-to-trial based on the strength of the last VTA DA neuron signal it experienced.

**Figure 2:**
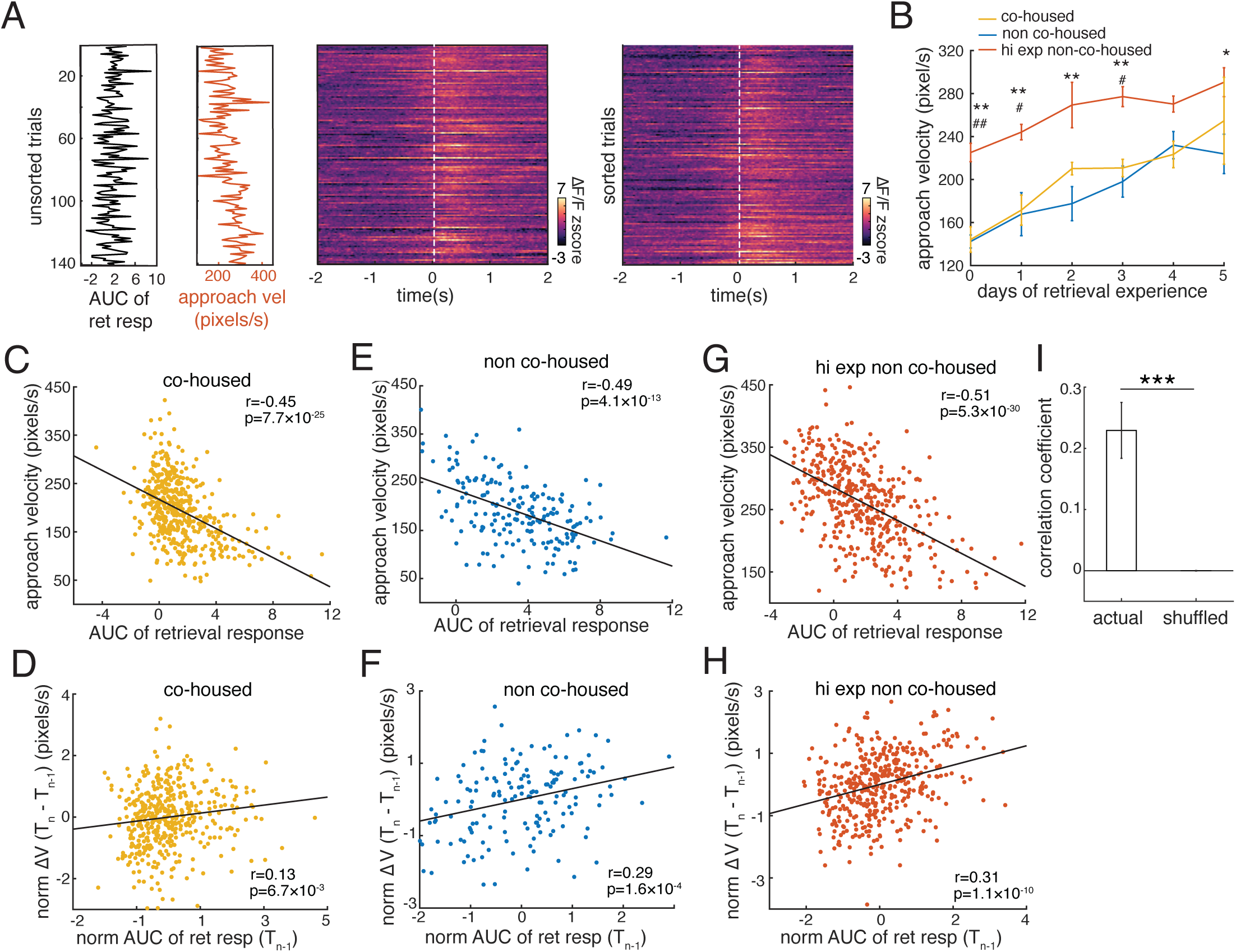
Retrieval behavior is updated from trial to trial based on VTA DA neuron signals. **(A)** Behavioral and neural data from one example mouse. *left*: Aligned vertical plots of the magnitude of VTA DA neuron activity (black trace) and approach velocity (red trace) for each of 140 trials over 6 days. r*ight*: Heat maps of neural activity for the same 140 trials. Each row is one trial. The left plot is sorted in chronological order and the right plot is sorted by approach velocity. **(B)** Plot of mean approach velocity over days for all three cohorts of mice depicted in Figure 1 (same groups as Fig 1, two-way ANOVA with Tukey correction, main factor (day): *p<0*.*0001*, main factor (mouse type): *p<0*.*0001*; * and # represent significant difference in the CH vs the HE/NCH groups, and in the NCH vs the HE/NCH groups respectively, \**p* < 0.05, \*\**p* < 0.01). **(C, D)** Scatterplots of approach velocity versus VTA signal amplitude (C) and change in approach velocity since the previous trial versus the VTA signal amplitude on the previous trial for the CH cohort. **(E, F)** Same plots as (C, D) but for the NCH cohort. **(G, H)** Same plots as (C, D) but for the HE/NCH cohort. **(I)** Bar plot of mean correlation coefficient for the data in (D), (F), and (H) comparing the result for matched trials versus shuffled trials (n = 13 mice, Wilcoxon matched pairs signed rank test, \*\*\**p<0*.*001*).

To test the effect of reward expectation on VTA DA neuron activity, and to observe the dynamics of responses to both pups and a cue signaling an imminent pup interaction, we devised a cued retrieval task (Fig. 3A and Methods). Briefly, the virgin mouse began each trial with an empty nest outside a two-chambered structure. A pup was placed in one of two chambers, either an inaccessible chamber or a chamber that could be accessed through a motorized sliding door that was closed between trials. On each trial, one of two tones was played, either a tone (CS+) that usually (>90%) signaled that the pup was in the accessible chamber, or a tone (CS-) that usually (>90%) signaled that the pup was in the inaccessible chamber (CS-). After a fixed delay (1 s from the end of the tone), the sliding door opened and the mouse was permitted to enter the chamber to search for a pup. Figure 3B shows mean VTA signals aligned to the tone, the door opening, and chamber entry, and separated into trials with an accessible pup (red) and trials without an accessible pup (black). Trials were randomized within a day and were carried out according to a standardized schedule on P0 – P8 (Fig. S3).

**Figure 3:**
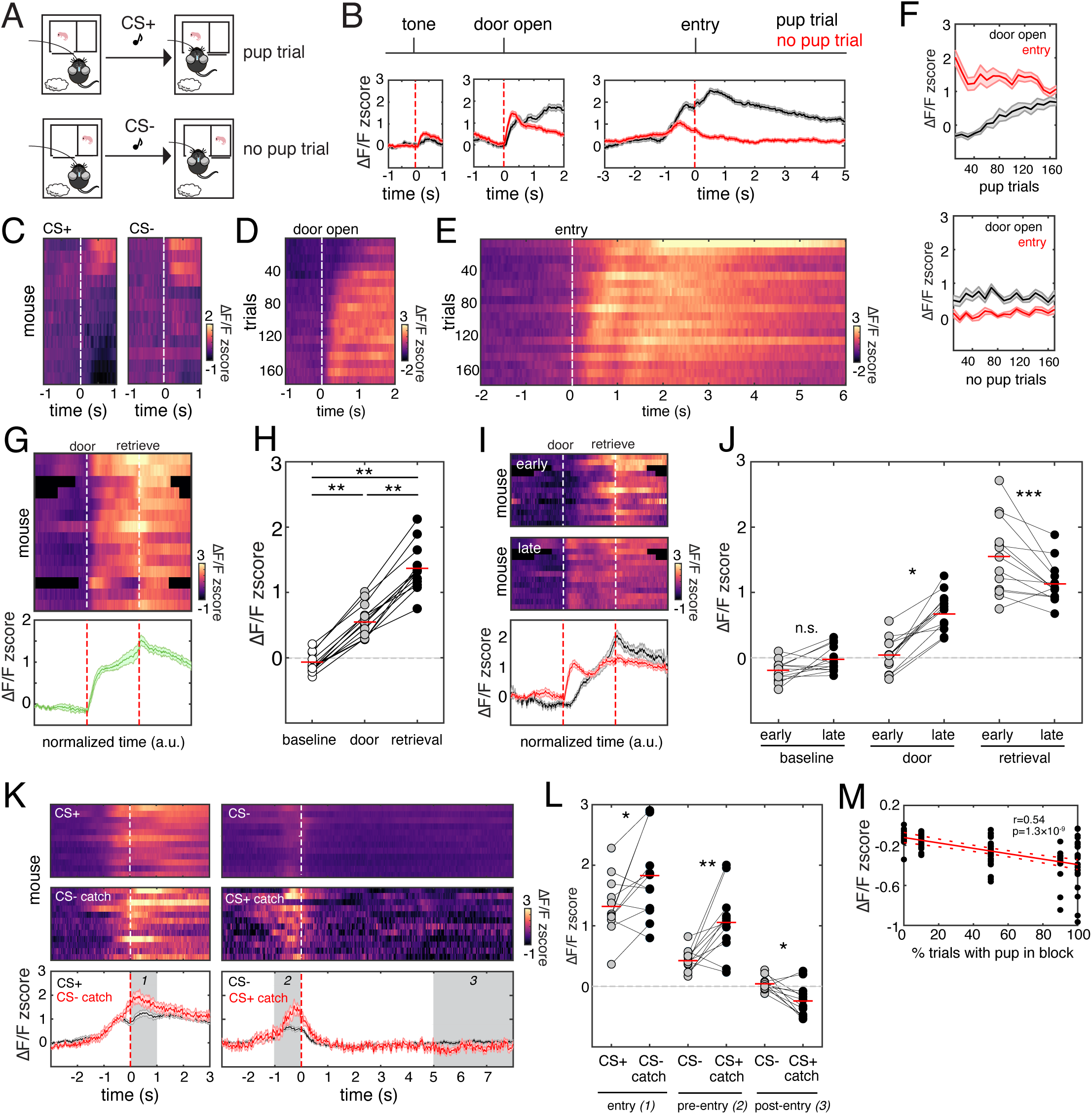
VTA DA neuron activity in a cued retrieval task is modulated by expectation. **(A)** Schematic of cued retrieval task. **(B)** Mean traces of VTA DA neural activity aligned to trial events for all mice comparing trials where the female encountered a pup (red) and trials where it did not (black). **(C)** Heatmap depicting the mean VTA dopamine neuron response to CS+ and CS-over the whole experiment for each mouse. **(D, E)** Heatmaps depicting the development of VTA dopamine neuron responses to door opening (D) and chamber entry (E) on CS+ trials averaged over all mice. Each row corresponds to the mean activity in bins of 10 trials. **(F)** Plots of mean ± SEM of the amplitude of responses to door opening and chamber entry over trials for all mice. The top plot depicts data from trials where a pup was encountered, and the bottom plot depicts data from trials where no pup was encountered. **(G)** Plot of data after time warping each trial and each mouse to align with both the door opening and retrieval. **(H)** Scatterplot of the mean signal for all mice comparing baseline and responses to the door and pup contact (n=13 mice, one-way ANOVA, all comparisons: *p* < 0.01) **(I)** *upper panels:* Heatmaps of the same data in (G) split into neural activity during the first 40 trials (‘early’) and the last 40 trials (‘late’) where a pup was encountered. *lower panel:* Plot of the mean activity during early trials (black) and late trials (red). **(J)** Scatterplot comparing the mean activity between early and late trials for baseline and responses to the door and pup contact. (n=13 mice, paired t test comparison with Bonferroni correction, *p* < 0.05) **(K)** *top:* Heatmaps depicting the mean activity for each of four trial types (CS+, CS+ catch, CS-, CS-catch). Each row is the mean VTA DA neuron activity for one mouse. *bottom:* Mean traces of each trial type comparing entry activity of (*left*) CS-catch trials (red trace) to CS+ trials (black trace) and (right) CS+ catch trials (red trace) to CS-trials (black trace). Gray shaded regions denote the data ranges used to calculate mean signals between normal and catch trials in (L). **(L)** Scatterplot comparing the mean DA neuron activity between normal and catch trials for each of the three windows marked in (K) (n=12 mice, paired t test comparison with Holm-Bonferroni correction, \**p* < 0.05, \*\**p* < 0.01) **(M)** Scatterplot with a linear correlation fit of mean baseline VTA DA neuron activity during each behavior block for all mice over the fraction of trials in which the mouse encountered a pup.

VTA DA neuron responses to task events changed over training. Most mice exhibited weak if any response to either tone (Fig. 3C). This was surprising because the CS+ predicted a rewarding pup encounter, however we did observe stronger and more consistent responses to the door opening, an event that was more proximal to pup contact (Fig. 3D). Considering trials where the female encountered and retrieved a pup, we plotted the mean activity for all mice in 10 trial bins (Fig. 3, D and E). As expected, there were large responses to chamber entry, which were followed quickly by pup contact in these trials. For trials where the female encountered a pup, the entry response grew weaker and the door response grew stronger (Fig. 3F). Activity was unchanged over trials without a pup for both door opening and chamber entry (Fig. 3F).

To more precisely quantify this shift in DA neuron responses over time from the pup to a pup-predictive stimulus, we used linear time warping to align the neural data to both the opening of the door and the retrieval of the pup (Fig. S4 and Methods). Aligning the data in this way revealed precise and distinct responses to the door and pup contact that were significantly different from baseline and each other (Fig. 3G and H). Comparing the magnitude of each peak between the first 40 pup trials (‘early’) and the last 40 pup trials (‘late’) showed a significant increase in responses to the door and a significant decrease in responses to pup contact in late trials (Fig. 3I and J). This pattern of VTA DA neuron activity strongly resembles changes in RPE over time.

To examine the effects of expectation on responses to task events, we manipulated the mouse’s prediction of access to a pup. The CS+ and CS-tones were not perfectly reliable cues for whether the pup would be accessible on a given trial. We included up to 10% ‘catch trials’ on which either the CS-tone was played and there was unexpectedly a pup in the chamber (CS-catch trials), or the CS+ tone was played and the pup was unexpectedly absent from the chamber (CS+ catch trials) (Fig. S3). Fig. 3, K and L compares the mean activity for all mice between catch trials and standard trials. The mean activity upon entry on CS-catch trials was significantly higher than on CS+ trials (Fig. 3, K and L). In both trials, the mouse experienced the same outcome, but on the catch trial, the presence of the pup was unexpected. In contrast, on CS+ catch trials, the activity just before entry was significantly greater than that on CS-trials, potentially in anticipation of pup contact (Fig. 3, K and L). Later in the trial, once the mouse had time to search the chamber and found no pup, there was a small, but significant dip in VTA activity (Fig. 3, K and L). This pattern of results is also consistent with VTA signals representing RPE for pup contact.

In addition to the amplitude of retrieval-related VTA activity being modulated by expectation, the baseline fluorescence reflected the recent trial history. We presented trials in daily blocks according to the schedule in Fig. S3. The percentage of trials where a pup was found in the accessible chamber varied from 0 – 100%. Across all mice, mean baseline fluorescence in a block was significantly inversely correlated with the proportion of trials where a pup was encountered in the chamber (Fig. 3M). This could be a mechanism for scaling the signal to noise ratio for reward responses to the statistics of pup encounters.

Finally, we tested whether VTA DA neuron activity is essential for rapid retrieval learning. DAT-Cre^+/−^ mice were injected in VTA of both hemispheres with a Cre-dependent AAV driving expression of either the inhibitory optogenetic tool stGtACR (*25*) or EGFP as a control (Fig. 4, A and B, S5). Naïve virgin females were tested with 20 daily trials of open field pup retrieval (P0 – P3). On each trial, a barrier was lifted, revealing a pup, and the mouse was free to retrieve it at any time during the trial (180 s) (Fig. 4C). We adopted a closed-loop design for triggering inactivation of VTA neurons with light on alternating trials, based on proximity to the pup, which is when VTA peaks. We used real-time tracking of the mouse’s head to trigger continuous light to the VTA while the mouse’s snout was in a circular zone around the pup’s starting location (6.6 cm dia.) (Fig. 4D). Prior to closed-loop behavior, all subjects were tested with a standard retrieval assay. There was no significant difference between the performance of stGtACR-expressing and EGFP-expressing mice on this pre-test (Fig. 4E). Pre-test performance was also not correlated with their performance during light inactivation trials on P0 (Fig. 4F).

**Figure 4:**
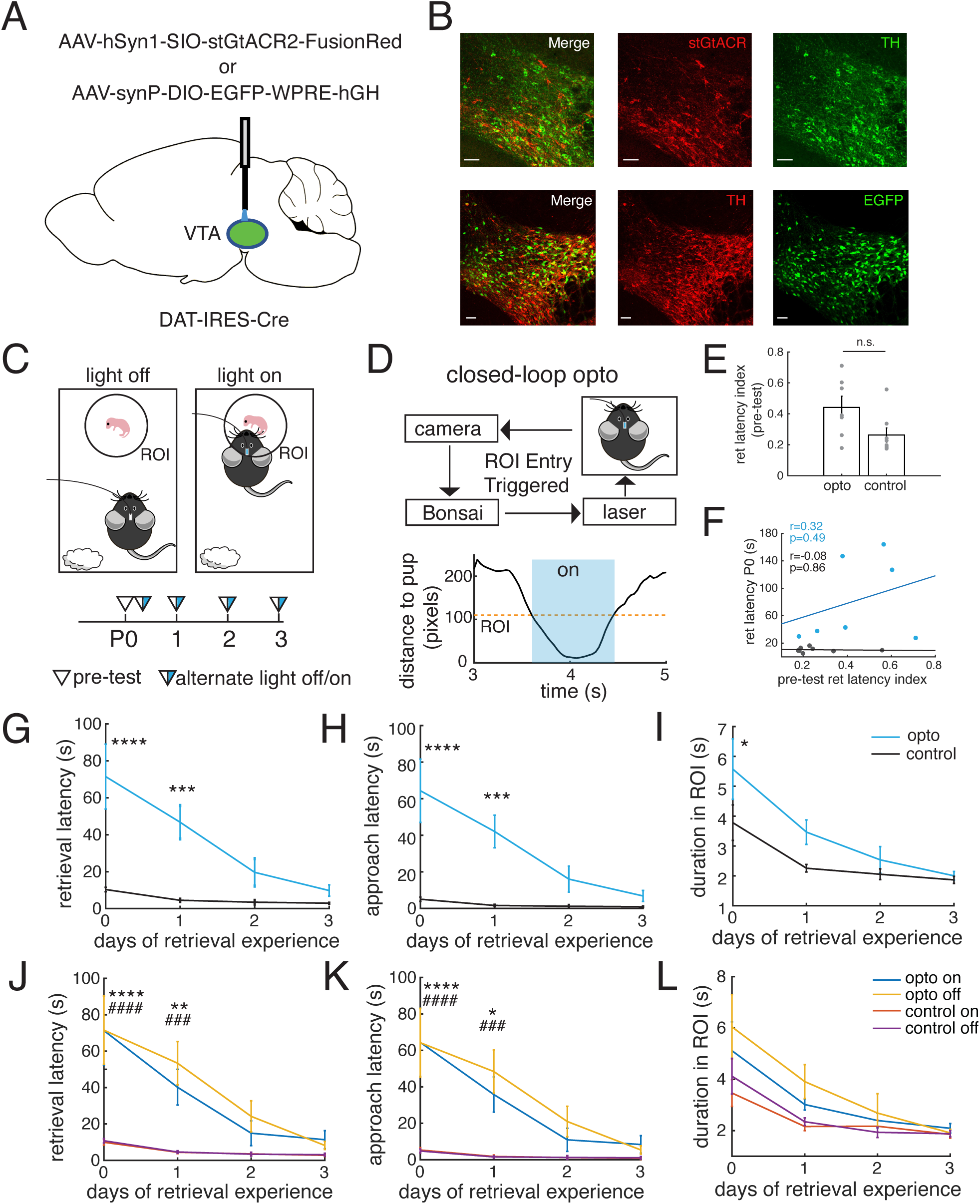
VTA DA neuron activity at pup contact gradually reinforces pup retrieval. **(A)** Intersectional viral strategy for optogenetic inhibition of VTA. **(B)** Co-localization of stGtACR (red) and TH (green), and EGFP (green) and TH (red) in VTA DA neurons. Scale bar=50 µm. **(C)** Experimental design. Naïve subjects were asked to retrieve pups one at a time (40 trials/day, P0 – P3). VTA neurons were inhibited by trains of 473 nm light on every other trial (Supplementary Movie 1). **(D)** Schematic of system to deliver closed-loop optogenetic inhibition when the mouse enters the ROI around the pup’s starting location. **(E)** A bar plot of data from a retrieval pre-test administered before the start of the experiment showed no significant difference in performance before training (n = 7 and 8 for stGtACR-expressing and EGFP-expressing mice, Mann-Whitney test, *p* > 0.05). **(F)** Scatterplot showing no relationship between pre-test performance and the effects of inactivating VTA DA neurons on P0. **(G – I)** Plots of retrieval latency (F) (two-way ANOVA with Sidak correction, main factor (day): *p* < 0.0001, main factor (mouse type): *p* < 0.001, *** *p* < 0.001, **** *p* < 0.0001), approach latency (G) (two-way ANOVA with Sidak correction, main factor (day): *p* < 0.0001, main factor (mouse type): *p* < 0.001, *** *p* < 0.001, **** *p* < 0.0001), and time in trigger zone (H) (two-way ANOVA with Sidak correction, main factor (day): *p* < 0.0001, main factor (mouse type): *p* < 0.05, * *p* < 0.05) across days comparing stGtACR-expressing (blue) and GFP-expressing (black) mice. **(J – L)** Same as (G – I) but with light on and light off trials plotted separately (two-way ANOVA with Tukey correction, main factor (day): *p* < 0.0001 for G-I, main factor (mouse type): *p* < 0.0001 for G-H and *p* > 0.05 for I, \*\**p* < 0.01, \*\*\**p* < 0.001, \*\*\*\**p* < 0.0001, * represents significant difference in opto on vs both control on and off groups, # represents significant difference in opto off vs both control on and off groups).

stGtACR-expressing mice were initially dramatically slower to return pups to the nest and took more trials to achieve the same performance when compared with EGFP-expressing controls (Fig. 4G, Supplementary Movie 2). This difference was predominantly due to delay in approaching the pup. stGtACR-expressing mice took significantly longer to approach the pup than EGFP-expressing mice (Fig. 4H, S6), but differed little in time spent in the trigger zone retrieving the pup (Fig. 4I, S6). Comparing trials with and without light inhibition, stGtACR-expressing mice exhibited indistinguishable overall retrieval latency, latency to approach, and time in the trigger zone (Fig. 4, I to K). Thus, light inhibition on a given trial was irrelevant to performance on that trial. Our interpretation of this result is that impaired retrieval performance reflects the cumulative effect of VTA inactivation on reward history, not acute interference with behavior.

Several neuromodulatory systems have been implicated in facilitating the emergence of maternal retrieval behavior including the oxytocin system (*26, 27*), noradrenaline system (*28, 29*), and dopamine system (*18–23*). Nevertheless, the mechanisms by which these systems orchestrate changes in social interaction remain unclear. Here we present evidence that maternal retrieval is established incrementally over many trials according to a common RL algorithm in which contact with the pup acts as the primary reward. We demonstrate that VTA DA neurons collectively signal RPE for this social reward and are essential for establishing and refining maternal retrieval. These findings complement recent work revealing that VTA DA neurons also play a role in other spontaneous natural behaviors including vocal imitation in juvenile songbirds (*30–32*) and response selection during social defeat (*33*).

More broadly, our results are consistent with a model of intraspecific interaction in which social decisions are cumulatively influenced by the motivation to maximize rewards obtained through social contact. Results in nonhuman primates reveal the quantifiably high value and fungibility of social stimuli in terms of non-social reward currency such as juice (*34, 35*). Therefore, this model may be a useful framework for understanding social decision making in diverse species, including humans.

## Acknowledgments

The authors would like to thank R. Mooney, S.R. Datta, A. Zador, B. Li, and members of the Shea Lab for helpful comments and discussion.

## Funding

National Institutes of Mental Health grant R01MH119250 (SDS)

The C.M. Robertson Foundation (SDS)

The Feil Foundation (SDS)

## Author contributions

Conceptualization: SDS, YX

Formal analysis: YX, SDS

Funding acquisition: SDS

Investigation: YX, AC, LH, AHP

Methodology: YX, LH, SDS

Software: LH, YX

Supervision: SDS

Visualization: YX, SDS

Writing – original draft: SDS, YX

Writing – review & editing: SDS, YX, LH, AC, AHP

## Competing interests

Authors declare that they have no competing interests.

## Data and materials availability

All data, code, and materials used in the analysis are available upon request

## Methods

### Animals

All procedures were approved by the Institutional Animal Care and Use Committee (IACUC) at Cold Spring Harbor Laboratory and performed in accordance with the National Institutes of Health’s Guide for the care and use of laboratory animals. Mice were maintained in a 12h light/12h dark cycle with *ad libitum* access to food and water. Test animals were adult DAT-Cre female mice over 8 weeks old (Slc6a3tm1.1(cre)Bkmn; The Jackson Laboratory, 006302). Pups and dams used in the experiments were from CBA/CaJ pairs. For optogenetics experiments, littermates were used for the control and the test groups.

### Genotyping

After weaning on P21, the offspring of DAT-Cre animals were genotyped by taking an ear punch. Each sample was first digested with 100 µL DirectPCR (Viagen, 402-E) and 1 µL Proteinase K (Invitrogen, 2036707) at 60 °C for 4 hours, and then the Proteinase K was inactivated at 95 °C for 45 minutes. The PCR solution for each sample contained 12.5 µL GoTaq Green Master Mix (Promega, M7123), 1.5 µL common forward primer, 1.5 µL reverse primer for the mutant or the wild-type allele, 2 µL genomic DNA, and 7.5 µL dH_2_O. Primer sequences, PCR reactions and electrophoresis were prepared according to a protocol from the Jackson Laboratory.

### Histology

Animals were deeply anesthetized with a lethal dose of Euthasol (Virbac, 200-071) via intraperitoneal injection, and were subsequently transcardially perfused with phosphate-buffered saline (PBS) and 4% paraformaldehyde (PFA; FD Neurotechnologies, PF101) at a flow rate of 5 mL/min. Brains were post-fixed in 4% PFA overnight at 4 °C and then transferred to a 30% sucrose solution in PBS. Brains were embedded with OCT compound (Sakura, 4583) and sectioned frozen at 50 µm on a sliding microtome (Leica, SM 2010R). For immunostaining, free-floating sections were washed with PBS 3 times and blocked in 5% normal goat serum (NGS, Vector Laboratories, S-1000-20) and 0.1% Triton X-100 (Sigma, T8787) at room temperature (RT) for 1 hour. Sections were incubated overnight with primary antibodies in a solution containing 0.5% NGS and 0.1% Triton X-100 at 4 °C. The next day, sections were incubated for 2 h with secondary antibodies in a solution containing 0.5% NGS and 0.1% Triton X-100 at room temperature. After washing 3 times with 1X PBS, sections were mounted on coverslips with Fluoromount-G (SouthernBiotech, 0100-01).

### Imaging and Image Analysis

Brain sections were imaged with a Zeiss LSM 710 confocal microscope. Sections that contained VTA were imaged and analyzed with Zeiss Zen software.

### Viruses

We used the following commercially-available viruses: AAV-syn-FLEX-jGCaMP7f-WPRE (2.8 ×10^13^ GC/mL, Addgene, 104492-AAV9), AAV-hSyn1-SIO-stGtACR2-FusionRed (1×10^13^ GC/mL, Addgene,105677-AAV1), AAV-synP-DIO-EGFP-WPRE-hGH (0.9×10^13^ GC/mL, Addgene,100043-AAV9). All viruses were aliquoted immediately on delivery and stored at −80 °C until use.

### Stereotaxic Surgery

Animals were injected intraperitoneally with a mixture of ketamine (100 mg/kg, Ketaset, NDC 54771-2013) and xylazine (5 mg/kg AnaSed, NDC 59399-110-20), and subcutaneously with meloxicam (1 – 2 mg/kg, Metacam) and Baytril (10 mg/kg, Bayer) subcutaneously prior to surgeries. Animals were placed in a stereotaxic device and maintained at a plane of anesthesia with 1 – 2% isoflurane mixed with oxygen at a flow rate of 1 – 2L/min. For fiber photometry experiments, 200 – 800nL Cre-dependent jGCaMP7f AAV was unilaterally injected into the VTA at 10 nL/min. Standard coordinates used for injections were as follows: AP −3.00 and ML +0.50 relative to bregma, and DV −4.10 relative to the brain surface) Optical fibers (200 µm dia., NA 0.39) from Thorlabs (CFMLC12U) were implanted above VTA (AP: −3.00, ML: +0.50, DV: −4.00 from brain surface) by slowly advancing at a speed of 1 mm/min. For optogenetics experiments, 120 – 200nL Cre-dependent stGtACR2 and EGFP AAV was bilaterally injected into the VTA (AP: −3.00, ML: −0.50 and +0.50, DV: −4.10 from brain surface). Dual fiber cannulae (200 µm dia., NA 0.37 from Doric Lenses, B280-2013_6) were implanted above VTA (AP: −3.00, ML: −0.50/+0.50, DV: −4.00 from brain surface). Real time tracking of the animal’s position during optogenetics experiments was achieved by minature infrared LEDs secured to the skull with dental cement (Metabond, C&B, S380). Animals were allowed to recover from surgery for 3 weeks before commencing experiments.

### Behavioral annotation and tracking

All behavior sessions were video recorded from above at 30 frames/s in a dark, sound-attenuated chamber under infrared illumination. Video data were acquired to a computer by either a Logitech webcam in fiber photometry experiments or an industrial camera (Teledyne FLIR, BlackFly S) in optogenetic experiments. Behavior sessions were conducted during the animals’ light phase (9:00h – 19:00h).

In pup retrieval behavior assays for fiber photometry experiments (Fig. 1 and 2), the test animals were habituated for 10 minutes in their home cages with a light and flexible optical fiber cable connected to their implant. We ran three cohorts of mice: (1) Co-housed (CH) surrogates were virgin DAT-Cre female mice that were introduced to the cage of pregnant, primiparous CBA/CaJ female 1 – 5 days before the dam gave birth. At that time, the sire was removed and the surrogate and dam were co-housed through P5. Each postnatal testing day (P0 – P5), the CH surrogate was tested alone with 5 scattered pups for 5 minutes in each of 3 sessions; (2) Non co-housed (NCH) surrogates were virgin DAT-Cre female mice that lived in their home cages with other virgin females only throughout the experiment. Each day from P0 through P5, the NCH surrogate was tested alone with 5 scattered pups for 5 minutes in 1 session; (3) High experience, non co-housed (HE/NCH) surrogates were virgin DAT-Cre female mice that each day from P0 through P5, the HE/NCH surrogates were tested alone with 8 scattered pups for 5 minutes for each of 3 sessions. Retrieval latency index was calculated as described (*36*).

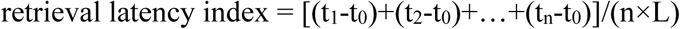

where n = number of pups scattered in the cage initially, t_0_ = time of the start of a trial, t_n_ = time of retrieving the nth pup, L = 300 s.

For the cued retrieval task with fiber photometry (Fig. 3), virgin DAT-Cre mice were co-housed with the CBA/CaJ dams through P8. Each day the surrogates were tested alone in their home cages. They were habituated for 5 – 10 minutes and then delivered snacks at random intervals to measure neural responses to unexpected snack. Subsequently, a chamber with a sliding door controlled by a stepper motor (McMaster-Carr, 6627T33), a stepper motor driver (McMaster-Carr, 6627T41), Arduino (Arduino, Uno R3) and Pulse Pal (Sanworks, v2) was placed in the home cage. Depending on the trial type, a pup was manually placed in the accessible or the inaccessible area inside the chamber. 40 trials were run each day except for P8 on which either 40 or 80 trials were run. Five seconds after a trial started, a CS+ (8 kHz tone) cue or a CS-cue (16 kHz tone) with a duration of 0.5 s or 2 s and a volume of 70 dB at animals’ heads was processed by an amplifier (Brownlee Precision, 410) and presented via an electrostatic speaker and its driver (TDT, ES1 and ED1). The door began to open 1 s after the tone stopped playing, and it took 2 s for the door to fully open. This allowed the animal to enter the chamber to search for a pup. The daily schedule of trials can be found in Fig. S3.

For optogenetics experiments (Fig. 4), virgin DAT-Cre mice were co-housed with pregnant CBA/CaJ dams until P0, and afterwards they were co-housed with their littermates. At the beginning of P0, the DAT-Cre surrogates were subjected to a pup retrieval assay (pre-test) for 5 minutes in which 5 pups were scattered in their home cages. Each day from P0 through P3, they were habituated with a dual optical fiber attached to their implant in their home cage for 10 minutes, followed by 20 alternating light off/on trials. In each trial, a pup was placed in the Region of Interest (ROI) for the test animal to retrieve.

Behaviors were annotated offline using the open source software package BORIS. Pup contact was defined as the first frame on which the animal put its mouth onto the pup at the end of approach in a pup retrieval trial. Retrieval was defined as the first frame on which the animals lifted the pup from the ground. The start of pup approach was defined as the moment animals walked out of the nest for pup retrieval. The end of pup approach was defined as the moment animals made contact with a pup. Snack eating was defined as the first frame that the animals bit on the snack in its hands.

To track animal position in the fiber photometry experiments, a deep learning-based method, DeepLabCut (*37*) was used. Traveled distance during approach in a pup retrieval trial was calculated based on the coordinates of the midpoint between the ears. Velocity was computed as the traveled distance of two consecutive frames multiplied by the frame rate, followed by convolution with a boxcar kernel of 7 points. Mean velocity of approach was calculated as the average of convoluted velocity during approach. For the videos in fiber photometry experiments, 1 pixel is equal to 0.07 cm. In optogenetics experiments, animal position was tracked in real time by a computer with a camera and running Bonsai (*38*) in which an image processing pipeline was used to extract the position of the LEDs on animals’ heads. For the videos in optogenetics experiments, 1 pixel is equal to 0.06 cm.

### Fiber Photometry

Fiber photometry was conducted as described (*28*). Briefly, the implant was coupled to a patch cord (Doric Lenses, P99414-01) through a mating sleeve (Thorlabs, ADAL1). Two LED light sources (470 nm and 565 nm; Thorlabs, M470F3 and M565F3;) were sinusoidally modulated by LED drivers (Thorlabs, LEDD1B) 180 degrees out of phase at a frequency of 211 Hz. Light power was adjusted to 30 µW at the start of each recording session. The emitted light passed through the patch cord, a focusing lens and dual edge dichroic mirrors to split it to separate photoreceivers (Newport, 2151). Signals were sampled at 6.1 kHz and acquired by NIDAQ boards (National Instruments, USB-6211 and USB-6001).

Data were analyzed with custom MATLAB (MathWorks) software. Briefly, the peaks of the two signals were extracted to achieve an effective sampling rate of 211 Hz and the data were filtered with a low-pass Butterworth filter with a corner frequency of 15 Hz. Signals were fitted with a double exponential function that was subtracted to correct for photobleaching. Next, we used a robust regression algorithm to fit a linear function for predicting the activity-independent component of the green fluorescence from the red fluorescence. The predicted green signal was subtracted and its mean was taken as baseline. ΔF/F was then calculated by dividing the residual green signal by the baseline.

The ΔF/F traces were converted to a Z-score using the mean and standard deviation of all signals recorded from a given subject. The Z-scored ΔF/F traces from individual trials were aligned to specific behaviors to generate heatmaps and compute mean fluorescence. When aligning the ΔF/F signals to specific behaviors, baseline fluctuations were removed by subtracting the mean signal in a time window extending 2 s prior to the event window. Responses to a given behavior during free retrieval (Fig. 1 and S1) were quantified as area under curve (AUC) for 4 s after the event. Responses to retrieval in Fig. 2 were quantified as AUC for −2 – 2s around the events. Responses to snack eating (Fig. S2) were measured as AUC for −4 – 4s around the events. To compare the changes of retrieval responses over days in different groups of animals (Figures 1), the AUC data were normalized to those on P0 within each animal. In the cued retrieval task, mean fluorescence was calculated for the designated window relative to the mean activity at the start of the trial before tone presentation. Linear time warping was used to align data to both the door opening and retrieval by aligning all traces to retrieval and then resampling to normalize each entire trial to the median time between the door opening and retrieval.

### Closed-loop optogenetics

The bilateral implant was coupled to an optical fiber that connected with a rotary joint (Doric Lenses, FRJ_1×2i_FC-2FC) and a 473-nm laser (OEM Laser Systems, PSU III LED). The laser driver was controlled by a Pulse Pal (Sanworks, v2) that received commands from Bonsai (*38*). The software also tracked the head position of the animal in real time with the aid of a miniature IR LED affixed to its head. Constant laser stimulation was delivered when the animal’s snout was detected inside the ROI before the pup was returned to the nest. The light power was adjusted to ~10 mW per hemisphere immediately before and after each experiment.

### In vivo extracellular electrophysiology

Virgin female DAT-Cre mice were injected unilaterally in VTA with AAV driving expression of Cre-dependent stGtACR as described above. After at least three weeks of recovery time, mice were acutely anesthetized with 1 – 2% isoflurane. Custom optrodes were constructed by gluing a 200 µm optical fiber to the shaft of a tungsten microelectrode (1.0 MΩ; Microprobe). Extracellular neural data were recorded with an AC differential amplifier (Model 1800; A–M Systems). Signals were bandpass filtered between 300 and 3000 Hz and digitally acquired to disk at 10 kHz with software and hardware from Cambridge Electronic Design (Power 1401, Spike2) for later spike sorting and analysis. Neurons encountered in the vicinity of VTA were tested with 5 – 15 10 mW pulses of constant 473 nm light (OEM Laser Systems) at 30 s intervals.

**Supplementary Figure 1:**
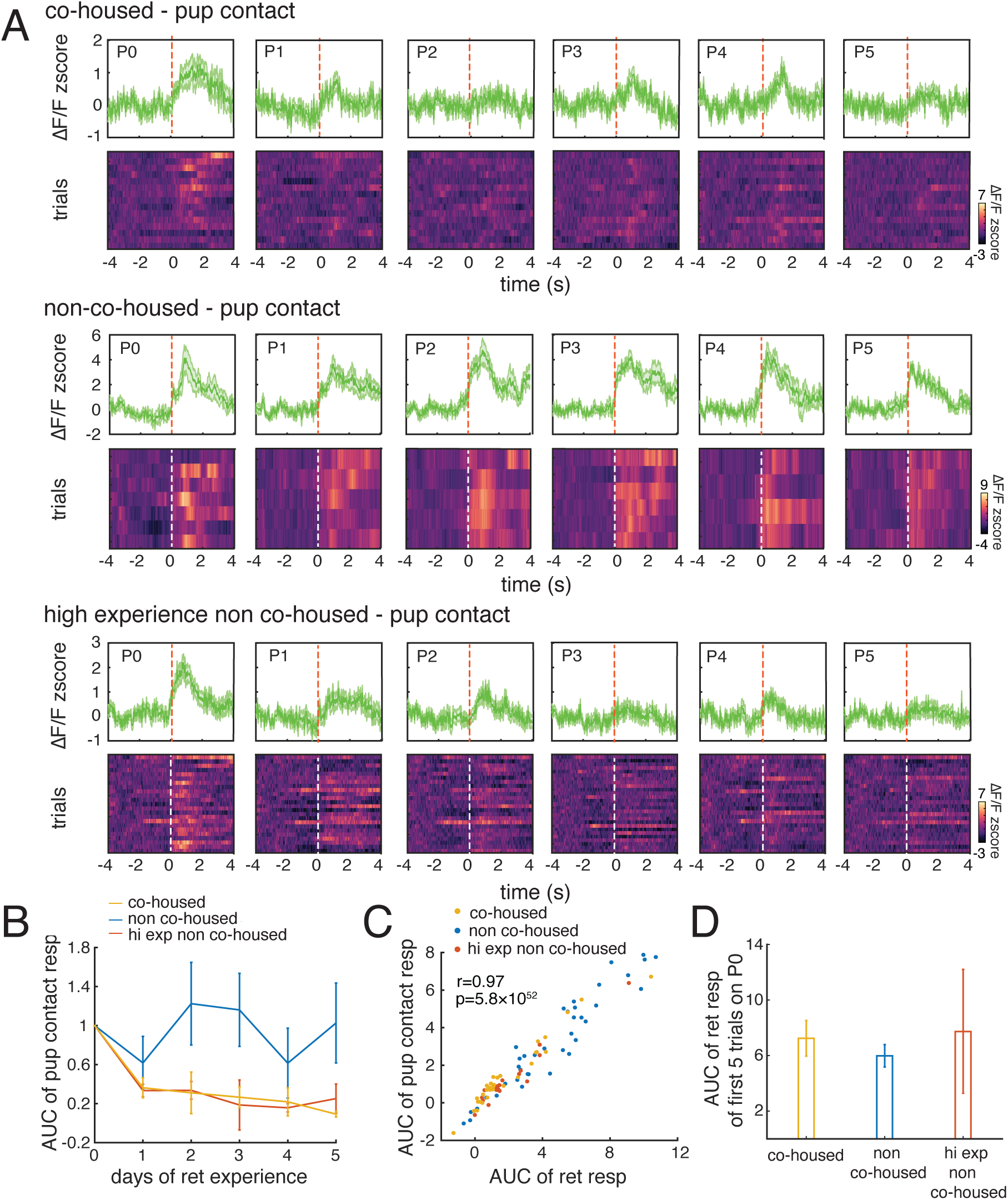
Comparison of data aligned to contact with data aligned to lifting the pup. **(A)** As in Figure 1D, but activity is aligned to the moment of contact with the pup. **(B)** As in Figure 1E, but the data are aligned to contact. (two-way ANOVA with Tukey correction, main factor (day): *p* < 0.01, main factor (mouse type): *p* < 0.01) **(C)** Scatterplot showing that the magnitude of the mean signal is nearly identical whether it is aligned to pup contact or pup lift. **(D)** Comparison of the mean VTA DA neuron signals on the first 5 trials for all three cohorts.

**Supplementary Figure 2:**
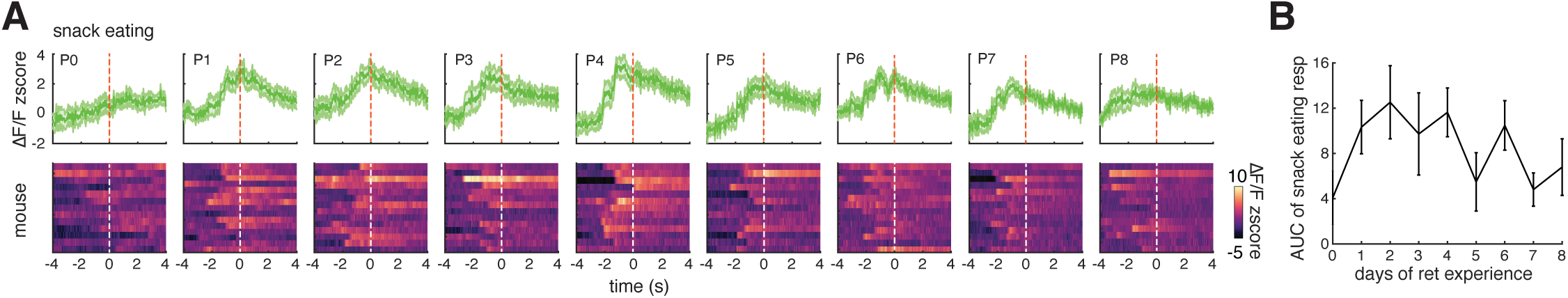
VTA DA neuron responses are of a similar magnitude to responses to an unexpected conventional reward. **(A)** Heatmaps (bottom row) and traces of mean +/− SEM (top) for responses of VTA DA neurons to eating an unexpected treat. Each row in the heatmaps is the response of one mouse for each day. **(B)** Mean integral amplitude of

**Supplementary Figure 3:**
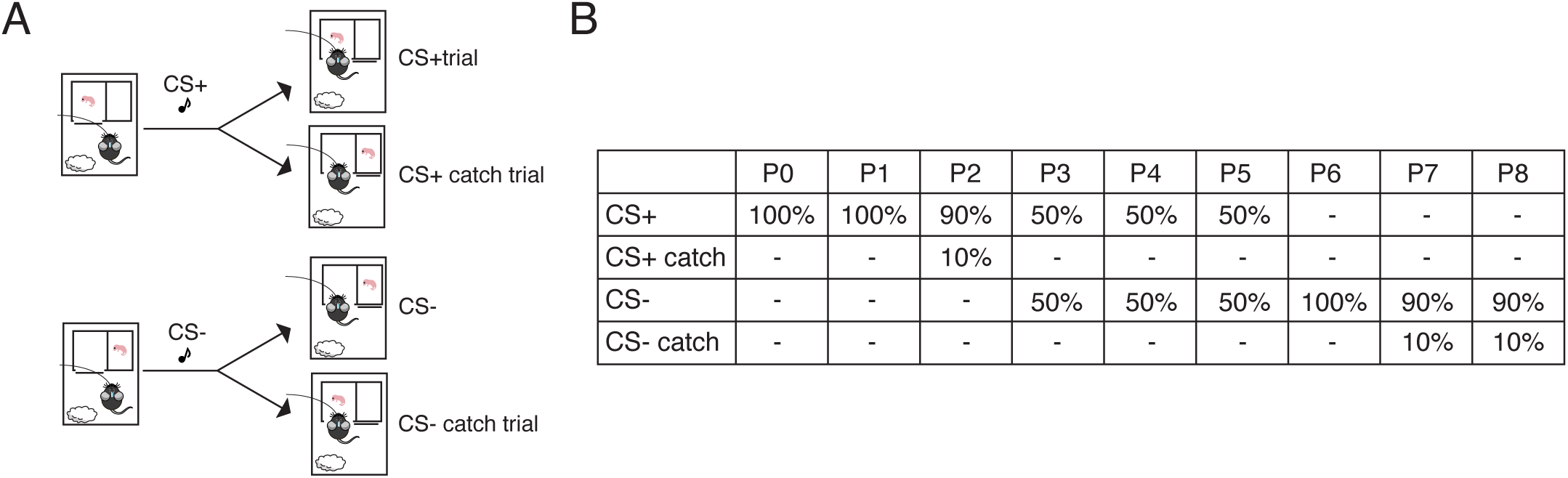
Cued retrieval behavior details. **(A)** Schematic of trial types. At the start of each trial, a mouse pup is placed in one of two chambers. One chamber is accessible through a sliding door and the other is not. The mouse is presented with one of two tones. One tone, designated CS+, indicates >90% of the time that the pup is in the accessible chamber. The other tone, designated CS-, indicates >90% of the time that the pup is in the inaccessible chamber. On some blocks, in 10% of CS+ trials, the pup is inaccessible, and in 10% of CS-trials, the pup is accessible. These are designated as CS+ catch trials and CS-catch trials, respectively. **(B)** Table indicating the daily behavior protocol each subject experienced from P0 - P8.

**Supplementary Figure 4:**
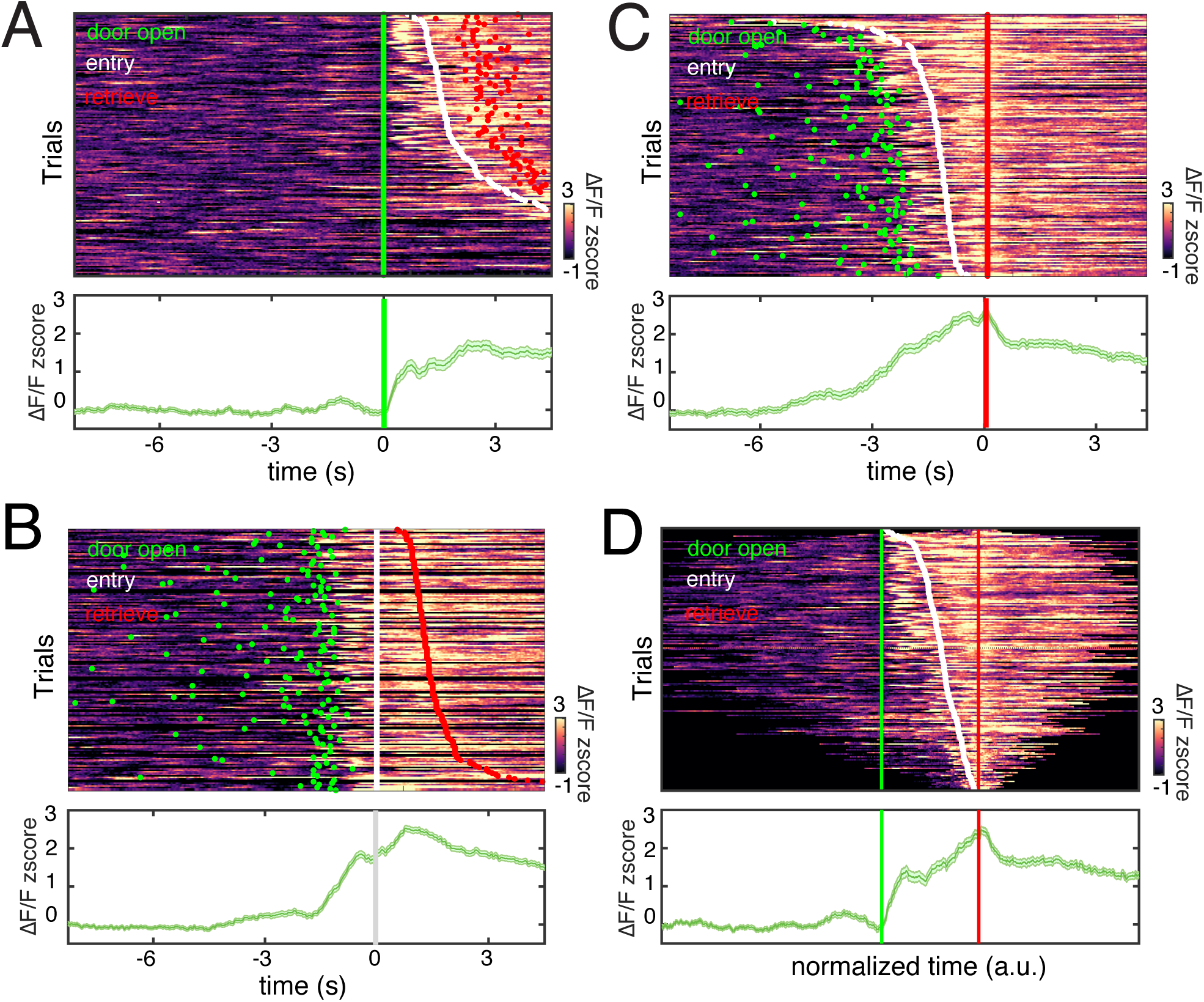
Alignment of trial events with VTA DA neuron signals. On individual trials, subjects exhibited variable timing between the opening of the sliding door (’door open’, green dots), stepping over the threshold of the chamber (’entry’, white dots), and contacting the pup (’retrieve’, red dots). We aligned the data to each of these events and also warped time to align the entire trial **(A)** Plot of VTA signals aligned to the opening of the sliding door. *upper panel:* Each row in the heatmap depicts one trial. The times of door opening, entry, and retrieval are indicated by the colored dots, and trials are sorted by the delay between the door opening and entry. The data represent 183 trials over 9 days from one mouse. *lower panel:* Plot of mean +/− S.E.M. of the neural activity in the upper panel **(B)** Plot of the same data aligned to retrieval and sorted by entry. Panels are otherwise organized as in (A). **(C)** Plot of the same data aligned to entry and sorted by retrieval. Panels are otherwise organized as in (A). **(D)** Plot of the same data with time stretched or compressed to standardize the interval between door opening and retrieval. Trials are sorted by entry time. Panels are otherwise organized as in (A).

**Supplementary Figure 5:**
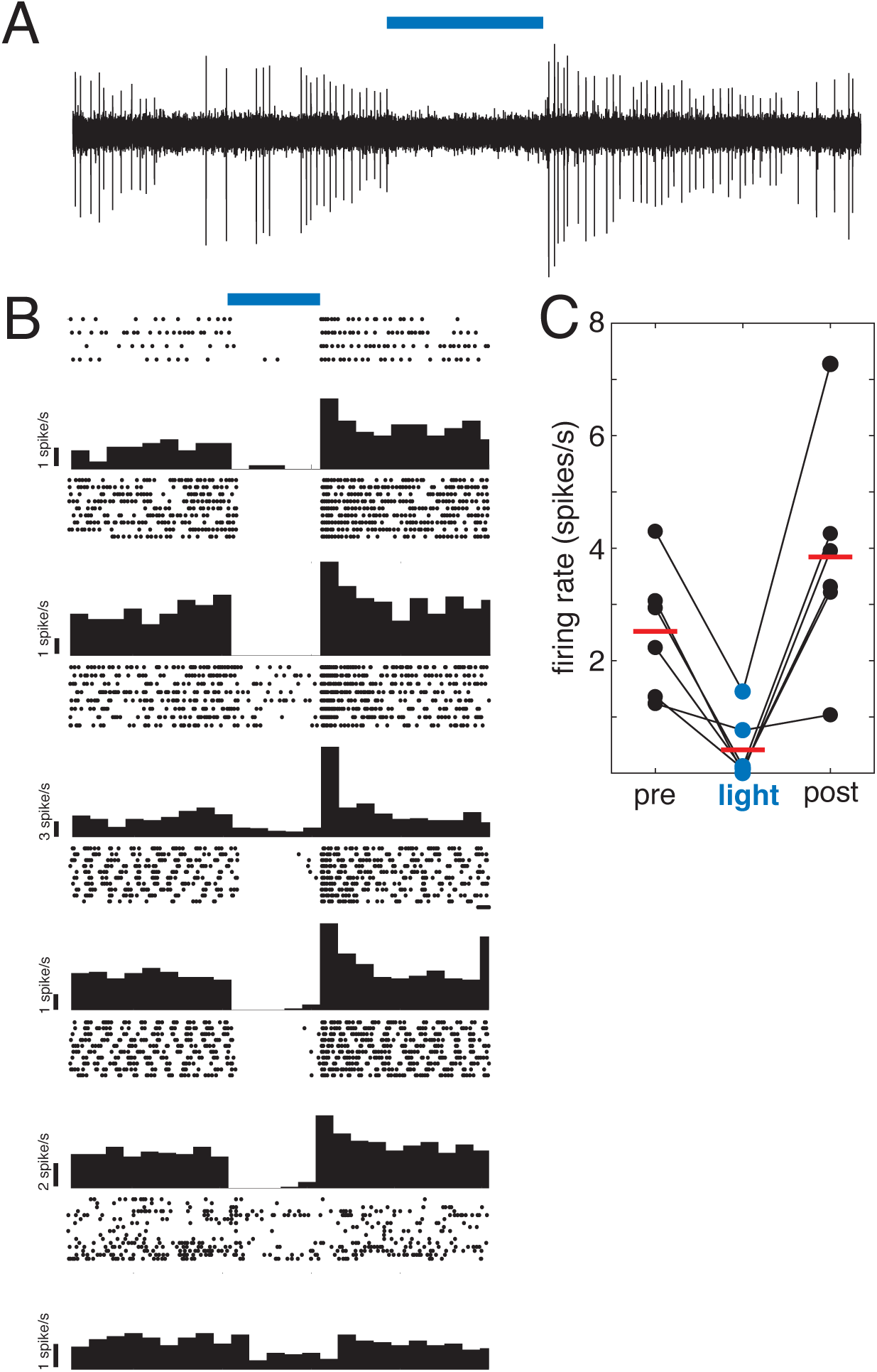
Fiber optic light delivery dramatically suppresses firing in VTA neurons expressing stGtACR *in vivo*. **(A)** Plot of a 25 s voltage trace from a recording of a single VTA neuron expressing stGtACR that includes a 5 s continuous 10 mW, 473 nm light pulse delivered by an optical fiber glued to the metal electrode. The light stimulus is denoted by the blue bar above the trace. **(B)** Raster plots and peristimulus time histograms summarizing the mean response to multiple light trials in 6 different neurons recorded in 2 mice. **(C)** Plot comparing the mean pre-light (5 s) firing rate, the mean light (5 s) firing rate, and the mean post-light (5 s) firing rate for the 6 neurons depicted in (B). The mean firing rate during the light delivery was significantly lower than both the mean pre- and post-light firing rates. One way ANOVA with Tukey’s HSD posthoc test. *p* < 0.01.

**Supplementary Figure 6:**
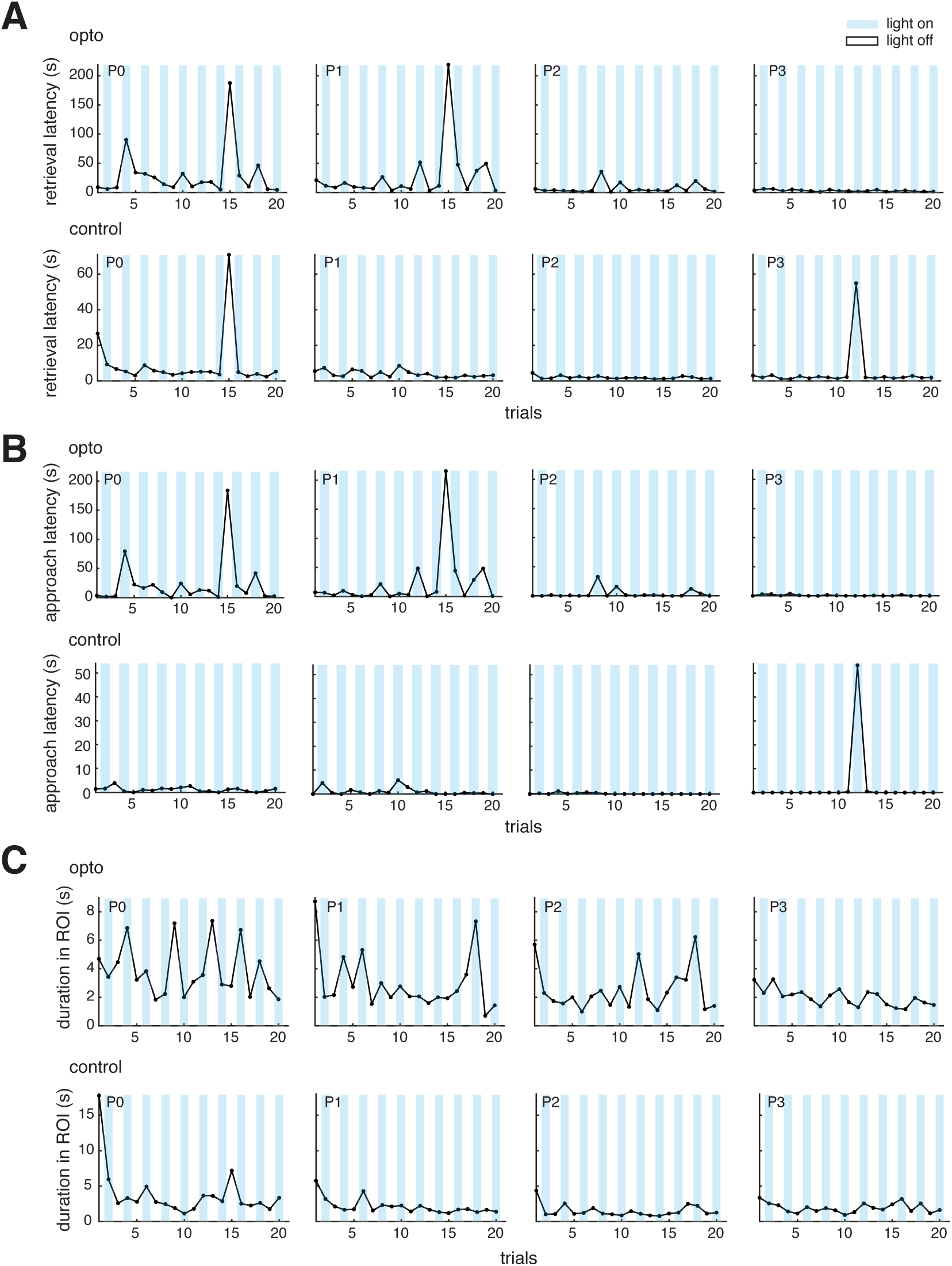
Comparison of trial latencies for an stGtACR-expressing mouse and a GFP-expressing control mouse. **(A)** Plots of total retrieval latency for all trials over four days (P0 - P3) from an optogenetically-inhibited mouse (top) and a control mouse (bottom) Trials with light delivery are denoted with a blue background. **(B)** Same as (A), but latency to approach the pup is plotted for each trial. **(C)** Same as (A), but time in the trigger zone is plotted for each trial.

## Notes

### Competing Interest Statement

The authors have declared no competing interest.

